# Immature HIV-1 assembles from Gag dimers leaving partial hexamers at lattice edges as substrates for proteolytic maturation

**DOI:** 10.1101/2020.10.01.322081

**Authors:** Aaron Tan, Alexander J. Pak, Dustin R. Morado, Gregory A. Voth, John A. G. Briggs

## Abstract

The CA (capsid) domain of immature HIV-1 Gag and the adjacent spacer peptide 1 (SP1) play a key role in viral assembly by forming a lattice of CA hexamers, which adapts to viral envelope curvature by incorporating small lattice defects and a large gap at the site of budding. This lattice is stabilized by intra- and inter-hexameric CA-CA interactions, which are important in regulating viral assembly and maturation. We applied subtomogram averaging and classification to determine the structure of CA at lattice edges and found that they form partial hexamers. These structures reveal the network of interactions formed by CA-SP1at the lattice edge. We also performed atomistic molecular dynamics simulations of CA-CA interactions stabilizing the immature lattice and of partial CA-SP1 helical bundles. Free energy calculations reveal increased propensity for helix-to-coil transitions in partial hexamers compared to complete six-helix bundles. Taken together, these results suggest that the CA dimer is the basic unit of lattice assembly, that partial hexamers exist at lattice edges, that these are in a helix-coil dynamic equilibrium and that partial helical bundles are more likely to unfold, representing potential sites for HIV-1 maturation initiation.

**Significance Statement:** HIV-1 particle assembly is driven by the viral Gag protein, which oligomerizes into an hexameric array on the inner surface of the viral envelope, forming a truncated spherical lattice containing large and small gaps. Gag is then cut by the viral protease, disassembles and rearranges to form the mature, infectious virus. Here, we present structures and molecular dynamics simulations of the edges of the immature Gag lattice. Our analysis shows that Gag dimers are the basic assembly unit of the HIV-1 particle, that lattice edges are partial hexamers, and that partial hexamers are prone to structural changes allowing protease to cut Gag. These findings provide insights into assembly of the immature virus, its structure, and how it disassembles during maturation.

## Introduction

The polyprotein Gag is the main structural component of HIV-1, consisting of the MA (matrix), CA (capsid), NC (nucleocapsid) and p6 domains as well as the spacer peptides SP1 and SP2 (1). Gag is produced in infected host cells and trafficked to the plasma membrane, where it assembles into a hexagonal lattice via its CA domain and recruits other viral proteins and the viral RNA genome (1, 2). Assembly of the curved Gag lattice is commensurate with membrane bending at the site of assembly, after which recruitment of Endosomal Sorting Complex Required for Transport III (ESCRT-III) components by the p6 domain of Gag induces membrane scission and release of the immature virus particle (2). The hexagonal Gag lattice accommodates curvature in the growing bud by incorporating vacancy defects (3). The activity of ESCRT-III is timed such that the final immature lattice is incomplete, giving rise to an additional large gap in the lattice, resulting in a truncated spherical shape (4, 5).

During or after budding, the viral protease is activated and cleaves this immature Gag lattice into its component domains, which leads to structural rearrangement within the virus particle (2). The released CA domains assemble to form a closed, conical capsid around the condensed ribonucleoprotein (RNP) complex of the mature virus (1, 6). Maturation is required for the virion to become infectious (1).

Within the immature virus particle, the N-terminal domain of CA (CA_NTD_) forms trimeric interactions linking three Gag hexamers while the C-terminal domain of CA (CA_CTD_) forms dimeric interactions mediated by helix 9 of CA, linking two Gag hexamers together (7). The CA_CTD_ additionally forms intra-hexamer interactions around the six-fold axis of the hexamer (7, 8). Amphipathic helices formed by the C-terminal residues of CA_CTD_ and the N-terminal residues of SP1 junction assemble into a six-helix bundle (6HB), thereby imposing hexagonal order on the CA domains, via classical knobs-in-holes packing mediated by exposed hydrophobic side chains, as also seen in coiled coils (8, 9). In combination, these relatively weak interactions give rise to a very dynamic, reversible assembly process that prevents the assembling lattice from becoming trapped in kinetically unfavorable states (10), as is the case with assembly of icosahedral viruses (11, 12). It is not surprising, therefore, that the energetics of Gag assembly are tightly controlled and highly dependent on scaffolding effects from the viral RNA and the membrane-interacting MA domain of Gag in order to ensure productive viral assembly (10, 13). Analysis of the diffusion pattern of fluorescently-labelled Gag supports the notion that Gag is trafficked to the site of assembly as low-order multimers, although it is still unclear whether these are Gag dimers, trimers or other multimeric forms of Gag (13, 14).

The primary assembly unit of the Gag lattice remains largely unknown. We can identify two hypothetical ways in which the lattice could assemble. First, the lattice could grow by addition of Gag hexamers (or sets of six component monomers), such that the CA-SP1 junction is assembled within a hexameric 6HB at all positions in the lattice. In this case interfaces between hexamers would be unoccupied at the edge of the lattice. From a purely energetic perspective, this appears most reasonable. Second, the lattice could form via addition of Gag dimers or Gag trimers (or equivalently from sets of either two or three component monomers). This would maintain, for example, the dimeric CA-CA inter-hexamer interactions but leave incomplete hexamers at the lattice edges, including unoccupied hexamer-forming interfaces along the CA-SP1 bundle. It additionally remains unclear whether the unoccupied Gag-Gag interfaces at the lattice edges are simply exposed, or whether they are stabilized by alternative conformations of individual domains or proteins, or by other binding partners. Understanding the structure of the edge of the immature Gag lattice therefore has implications for understanding the mechanism of virus assembly.

Viral assembly, budding and maturation are tightly linked and disrupting the kinetics of any of these processes can give rise to defects in maturation and formation of non-infectious viral particles (1, 15, 16). The rate-limiting proteolytic cleavage site in the maturation process resides within the CA-SP1 6HB (17). Unfolding of the helical bundle is required to allow proteolytic cleavage to proceed (18–20), but the exact mechanism for protease access to this site is not known. The spatial localization of proteolytic processing within the context of the immature Gag lattice is relevant: does the protease act on Gag within the lattice, or does it act on the edges of the Gag lattice, causing a cascade of lattice disruption? At the lattice edge, is the substrate for the protease with a 6HB or within an incomplete hexamer? Understanding the structure of the edge of the immature Gag lattice therefore has implications for understanding the mechanism of virus maturation.

High resolution immature Gag structures have previously been determined directly from purified viruses by cryo-electron tomography (cryo-ET) and subtomogram averaging (9). These structures represent an average hexamer within the immature lattice, with a full complement of 6 Gag hexamer neighbors. Here, we have applied subtomogram classification and averaging approaches to an existing immature virus dataset (9) in order to determine the structures of Gag assemblies at lattice edges. We also applied atomistic molecular dynamics simulations to assess the roles of the different CA-CA interactions in immature lattice stabilization, and to predict the properties of the structures we observe at lattice edges. Together, our results suggest that the basic unit of immature HIV-1assembly is a Gag dimer and that partial CA-SP1 helical bundles are present at the edges of the assembled lattice and may be substrates for initiation of maturation.

## Results

### Cryo-ET to reveal the structure of Gag at the lattice edge

As a starting point for analysis of the edges of the immature lattice, we took the cryo-ET dataset from which F. K. M. Schur et al. (9) previously determined a 4.2 Å map of immature HIV-1 CA and CA-SP1 directly from purified viruses (EMDB accession code: EMD-4017) (**Fig. 1A**). The data was partially reprocessed to ensure that as much of the Gag layer was retained in the dataset as possible (**Fig. 1B**). The coordinates of complete or partial Gag hexamers were computationally analyzed to identify those in the vicinity of lattice edges (**Fig. 1B**), which were subsequently used as input for further image classification.

**Figure 1.**
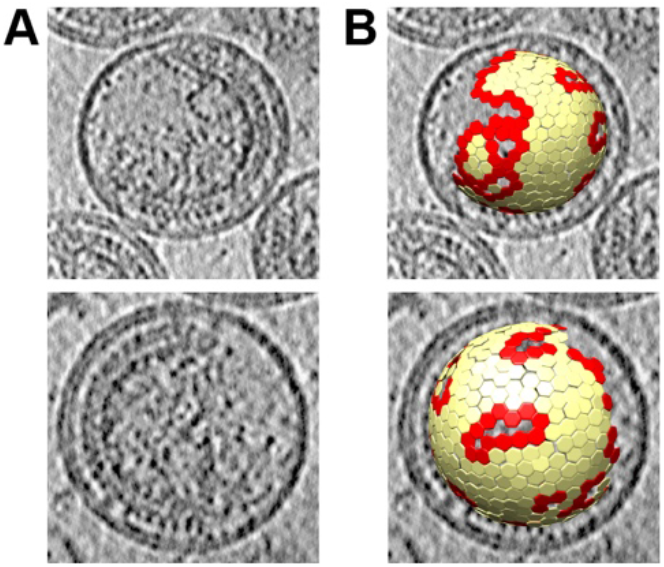
Illustration of immature HIV-1 virus particles and identification of Gag lattice edges. (A) Computational slices of 5.4 Å thickness through two representative tomograms from the dataset, illustrating the morphology of immature HIV-1 Gag lattice. An ordered Gag lattice is seen on one side of the virus with a large gap in the Gag lattice on the other side. (B) Lattice map showing aligned subtomogram positions corresponding to immature Gag hexamers, overlaid on the tomogram. Edge hexamers, defined as those with fewer than 6 hexamer neighbors, are shown in red and all other hexamers are colored in yellow.

Image classification of subtomograms aims to sort them based upon differences in macromolecular structure, but is complicated by noise in the data, and by the missing wedge problem (21–23): missing information in Fourier space due to physical limitations on the angular range across which a sample can be tilted in the electron microscope. Computational methods are required to compensate for this missing information (22). We employed two different classification approaches to achieve good separation of structural classes and to validate our results. These approaches were: 1) wedge-masked difference principal component analysis (WMD PCA) (23), and 2) multi-reference alignment and classification using synthetic references (24). These classification approaches are described in more detail in Materials and Methods and in Supplementary Fig. 1. Both classification approaches sorted the immature hexamers at the edge of the lattice, according to whether 1, 2 or 3 neighboring hexamers were missing. We did not identify hexamers lacking four or five neighboring hexamers, which could imply either that hexamer species lacking four or five neighbors do not exist at the edge of the lattice, or that these species exist but are excluded from the dataset because they do not align to a hexameric reference. Class membership differed between the two classification approaches when applied to the same input data, but this is not unusual for classification of noisy, missing wedge-affected data. Both approaches converged to similar structural classes (**Fig. 2, Supplementary Fig. 2, Supplementary Fig. 3**). These structural classes illustrate the variety of structures present at the lattice edge.

**Figure 2.**
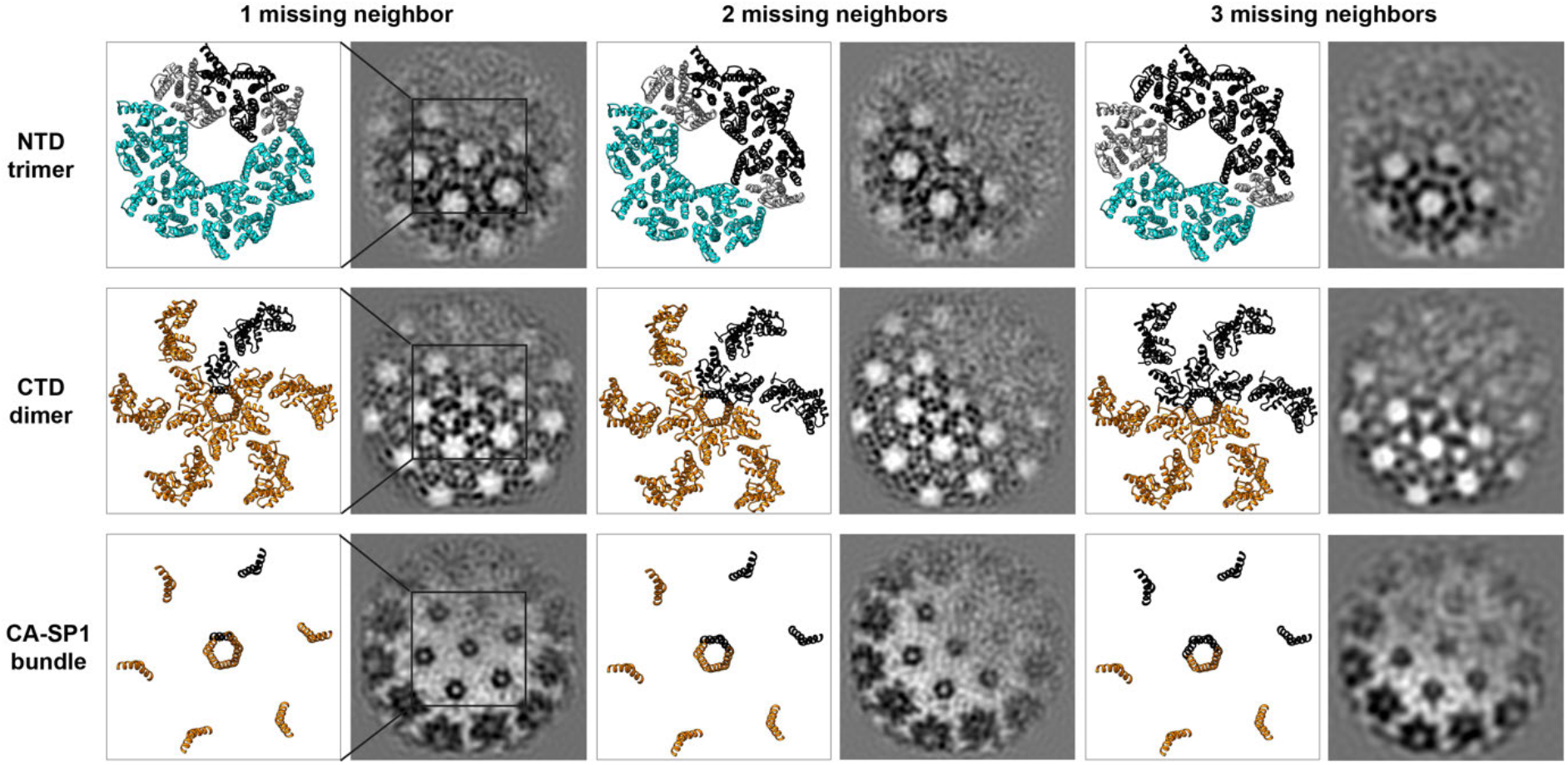
Cryo-EM structures obtained by WMD PCA classification of lattice edge hexamers. Classes with varying numbers of missing neighbors are shown next to corresponding CA_CTD_ and CA_NTD_ atomic models (PDB 5L93) fit as rigid bodies, with a box indicating the region of each class illustrated as an atomic model in each row. Models include the six monomers in the central hexamer, and the 18 surrounding monomers that interact with the central hexamer by either CA_CTD_ dimer or CA_NTD_ trimer interactions. Missing Gag molecules are depicted in black. CA_NTD_ trimers and CA_CTD_ dimers in which all trimer or dimer partners are present are shown in cyan and orange respectively. CA_NTD_ trimer positions which are missing trimer or dimer binding partners are shown in gray.

When we analyzed the appearance of the CA_CTD_ region of hexamers for which one neighboring hexamer was missing, we found that they were missing one CA_CTD_. Similarly, hexamers for which two or three neighboring hexamers were missing lacked two or three CA_CTD_s, (only four or three copies of CA_CTD_ were visible). Gag therefore appears to be behaving as a dimer – when one CA_CTD_ is absent, its dimeric partner is also absent. Note that hexamers lacking Gag subunits do not appear to relax into multimers with higher symmetry, e.g., hexamers missing one Gag subunit are not equivalent to Gag pentamers.

When one CA_CTD_ dimer is absent, the symmetry of the lattice means that one CA_NTD_ will be missing from each of two CA_NTD_ trimers. We observed that when one CA_CTD_ dimer is missing from a hexamer, the density for two CA_NTD_ trimers is missing. These data imply that when a CA_NTD_ trimer is lacking one member, the remaining two CA_NTD_ are no longer stabilized in their positions in the lattice and become mobile, hence they are not resolved (**Fig. 2**).

Together, these data imply that CA dimers are the basic assembly unit of the Gag lattice, and that the edge of the Gag lattice therefore consists of partial hexamers assembled from Gag dimers.

### Molecular dynamics of lattice edges

We performed atomistic molecular dynamics simulations to assess the impact of the dimer and trimer interfaces on the structural stability of CA-SP1 hexamers. We characterized stability by assessing the root mean squared deviation (RMSD) from the atomic model (PDB 5L93) at Cα resolution within each of the twelve α-helices (we denote the CA-SP1 junction as helix 12 or H12). A larger RMSD value qualitatively indicates a greater mean shift from the atomic model while a larger distribution of RMSD values suggests more structural variability; the median and interquartile range (IQR) of the RMSD are distinct yet related indicators of disorder. Hereafter, we will define a protein segment having an increasingly large median and IQR as being “more disordered.” As a baseline, the RMSD of a complete Gag hexamer (**Fig. 3A**) exhibited a median (IQR) of 3.3 (1.1) Å per helix.

**Figure 3.**
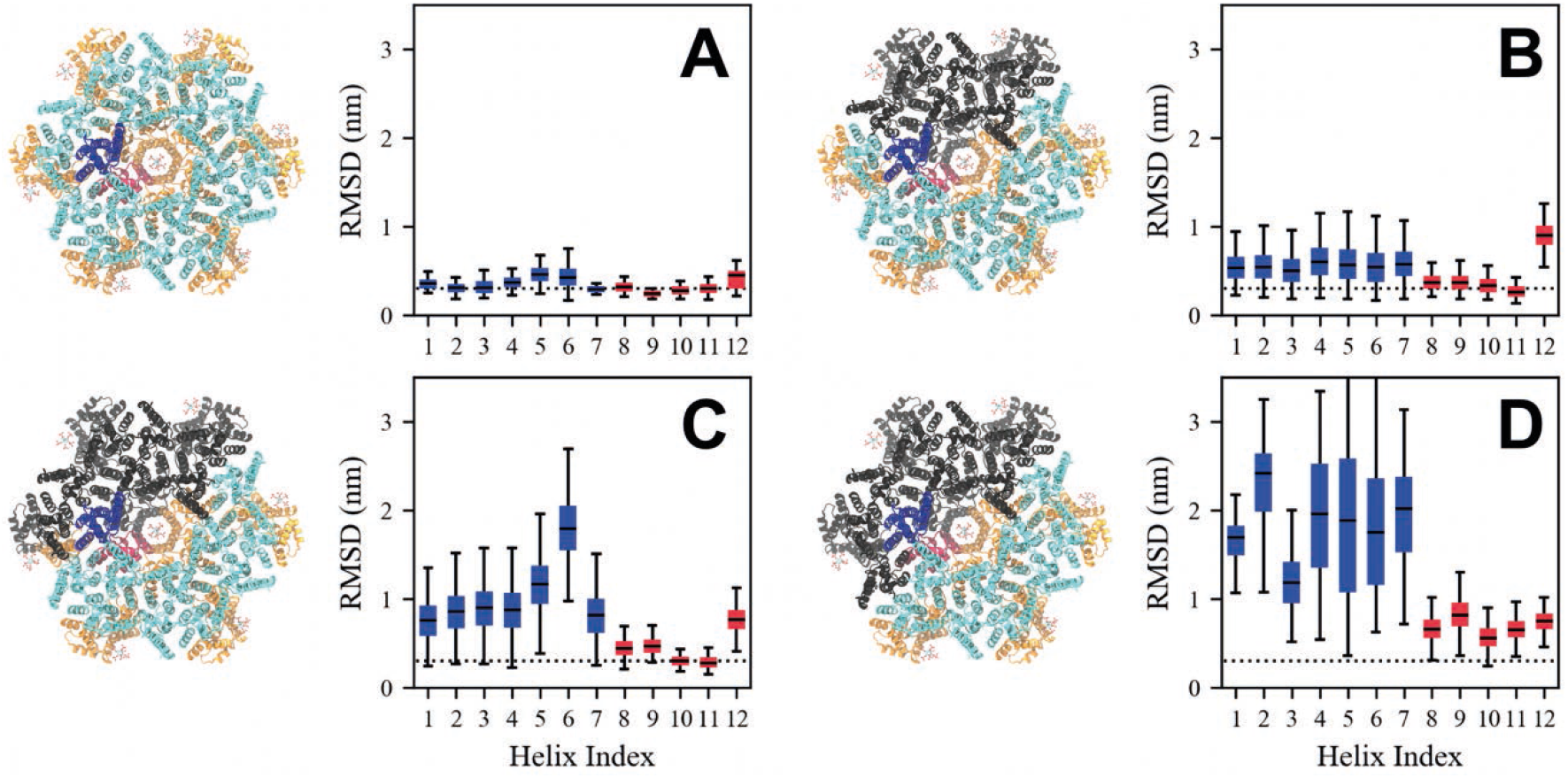
Comparison of the root mean squared deviation (RMSD) with respect to an atomic model (PDB 5L93) of a CA-SP1 monomer within (A) a complete hexamer and (B-D) incomplete hexamers missing 2 Gag subunits. The CA_NTD_ (CA_CTD_) of the analyzed monomer is colored blue (red), while the remaining CA_NTD_ (CA_CTD_) domains are colored cyan (orange). From (B-D), a Gag subunit adjacent to the analyzed monomer is removed such that the analyzed monomer maintains its CA_CTD_ dimer contact and one out of two CA_NTD_ trimer contacts, (C) maintains its CA_CTD_ dimer contact and no CA_NTD_ trimer contacts, and (D) lacks both CA_CTD_ dimer and CA_NTD_ trimer contacts; monomers that are missing are depicted in black. We note that the states analyzed in (A-C) are observed by cryo-EM while (D) is not and serves as a basis for comparison. Each box bounds the upper and lower quartiles with the central line indicating the median, while the whiskers show the extrema of the distributions. Blue (red) boxes refer to the analyzed CA_NTD_ (CA_CTD_) monomer. The dotted line marks a RMSD of 0.3 nm and serves as a guide to the eye.

We next considered an incomplete hexamer with 2 neighbors missing as observed in our cryo-EM dataset. In the clockwise-most CA-SP1 monomer, the CA_CTD_ is dimerized, while the CA_NTD_ has one, rather than two, of its trimer contacts (**Fig. 3B**). This CA-SP1 monomer has a median (IQR) of around 3.7 (1.2) Å per helix in the CA_CTD_ (helices 8-11), which suggests that the CA_CTD_ structure is similar to that of the complete CA-SP1 hexamer. The median (IQR) of the CA_NTD_ (helices 1-7), however, increases to around 5.4 (2.9) Å per helix, consistent with increasing disorder throughout the CA_NTD_ due to the absence of one trimer contact. Interestingly, removal of the second CA_NTD_ trimer binding partner (**Fig. 3C**) results in a further shift of the median (IQR) of the CA_NTD_ to around 17.9 (4.6) Å (most evident for H5 and H6) while the median (IQR) of the CA_CTD_ domain persists around 4.6 (1.2) Å per helix. Removal of the CA_CTD_ dimerization partner causes a significant increase in the median (IQR) to 17.5 (11.6) Å and 8.2 (2.4) Å per helix for both CA_NTD_ and CA_CTD_, respectively (**Fig. 3D**).

Taken together, our observations show that the loss of CA_NTD_ trimer contacts induces more disorder throughout the CA_NTD_ while the loss of CA_CTD_ dimer contacts induces more disorder in both the CA_NTD_ and CA_CTD_. These findings are consistent with the importance of the CA_CTD_ dimerization interface, and to a lesser extent, the CA_NTD_ trimerization interface, in stabilizing immature CA-SP1 hexamers. Moreover, our computational analysis supports our cryo-EM observations, which suggest that CA dimers act as the basic assembly unit of the Gag lattice.

### Cryo-ET suggests structured CA-SP1 regions in incomplete hexamers

In our density maps of incomplete lattice edge hexamers, we observe that when one or more Gag subunits are missing from a hexamer, the CA-SP1 6HB density becomes slightly weaker but does not disappear from the maps (**Figs. 2, 4**). The resolution of the maps is insufficient to characterize the bundle structure in hexamers missing Gag subunits, but the size and position of the density suggest that helical secondary structure is being maintained – once uncoiled, this region would not be expected to give rise to significant density (**Fig. 2**).

**Figure 4.**
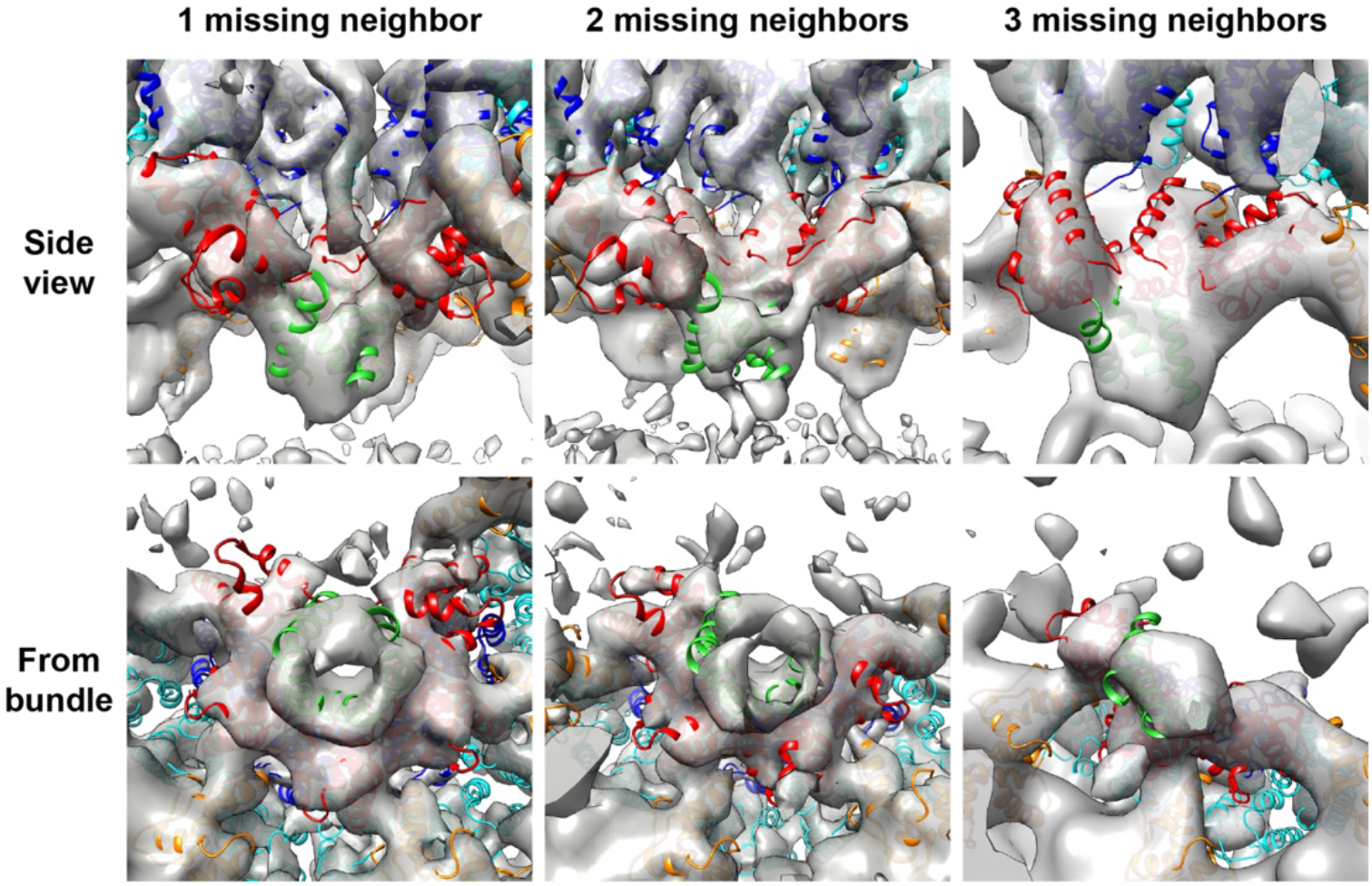
Side and top views of the CA-SP1 helical bundle region in the hexamer structures determined in lattice positions with 1, 2 or 3 missing neighboring hexamers. The CA_NTD_ and CA_CTD_ of the central, partial hexamer are depicted in blue and red respectively, whereas the CA_NTD_ and CA_CTD_ of neighboring hexamers are shown in cyan and orange respectively. The CA-SP1 helix in the partial bundles seen are shown in green.

Additionally, the density for the loop between H10 and H11 in the CA_CTD_ was observed to be weaker when the neighboring CA_CTD_ in the hexamer was absent, suggesting increased flexibility in the part of the CA_CTD_ directly upstream of a partial helical bundle (**Fig. 4**).

The presence of density in the CA-SP1 6HB region suggests that it can still exist in a stable form even when fewer than 6 helices are present and that the bundle structure can adapt to loss of a helix without becoming completely disordered. This raises the question of how the bundle accommodates loss of up to half of its constituent helices while retaining some ordered packing, given that a crystal structure of this region in a full CA-SP1 6HB exhibits classical knobs-in-holes packing of the hydrophobic residues exposed along the amphipathic CA-SP1 helix (8).

### Molecular dynamics to assess helix-coil transitions in the 6HB

Molecular dynamics simulations provided further insight into the structure and thermodynamics of the CA-SP1 6HB in both complete and incomplete hexamers. In all three incomplete hexamer cases studied above, we find that despite a median RMSD of around 8.0 Å for H12 (the CA-SP1 junction), the IQR remains low at 1.6 Å (**Fig. 3B-D**). Within our simulated timescale (around 410 ns), H12 maintains an α-helical secondary structure but tends to be distorted with respect to the 6HB quaternary structure. The same H12 in complete hexamers, however, has a small median and IQR of around 4.3 and 1.6 Å, respectively, and maintains both its α-helical and quaternary structure. These two observations suggest that the loss of neighboring CA-SP1 monomers distorts the quaternary structure of the 6HB while the secondary structure is maintained. However, it is known from nuclear magnetic resonance (NMR) spectroscopy experiments that the CA-SP1 region exists in a helix-coil equilibrium, even within complete hexamers (25). To assess the free energy of the helix-coil transition in both complete and incomplete hexamers, we performed Well-Tempered Metadynamics (WT-MetaD) simulations (see details in Methods).

We projected the free energy onto two coordinates. The first is the alpha-beta similarity (AB_sim_), which quantifies the number of phi and psi dihedral angles (*ϕ*) throughout H12 that are consistent with an α-helix; when the CA-SP1 junction is completely helical (or nonhelical), AB_sim_ is 30 (or 0). The second is the first component (tIC) from time-structure independent component analysis (tICA), which is a dimensional reduction technique used to identify the slowest varying linear projections of data. By construction, the first tIC refers to the slowest collective mode as described by changes to *ϕ*. Formal definitions and details on both of these coordinates can be found in the Methods section.

We compare the 2D-projected free energy surfaces for a helix in a complete hexamer, a helix with two neighboring helices in an incomplete hexamer, and the clockwise-most helix (i.e., an exposed helix with one neighbor) in an incomplete hexamer in **Fig. 5A-C**, respectively. The free energy surfaces for a helix with two adjacent helices in complete (**Fig. 5A**) and incomplete (**Fig. 5B**) hexamers appear to be qualitatively similar. The minimum free energy paths shown in **Fig. 5A-B** indicate that the free energy barrier heights for the helix-to-coil transition are comparable (8.5 and 9.0 kcal/mol, respectively). The free energy surface for the clockwise-most helix in the incomplete hexamer (**Fig. 5C**), on the other hand, is notably different. In particular, the free energy surface exhibits lower barrier heights, such as the 4 kcal/mol helix-to-coil barrier seen in its minimum free energy path (**Fig. 5C**). We also note that this helix appears to undergo a helix-to-coil transition following a different structural route, as indicated by the positional differences of the minimum free energy path in configurational space with respect to the former two minimum free energy paths.

**Figure 5.**
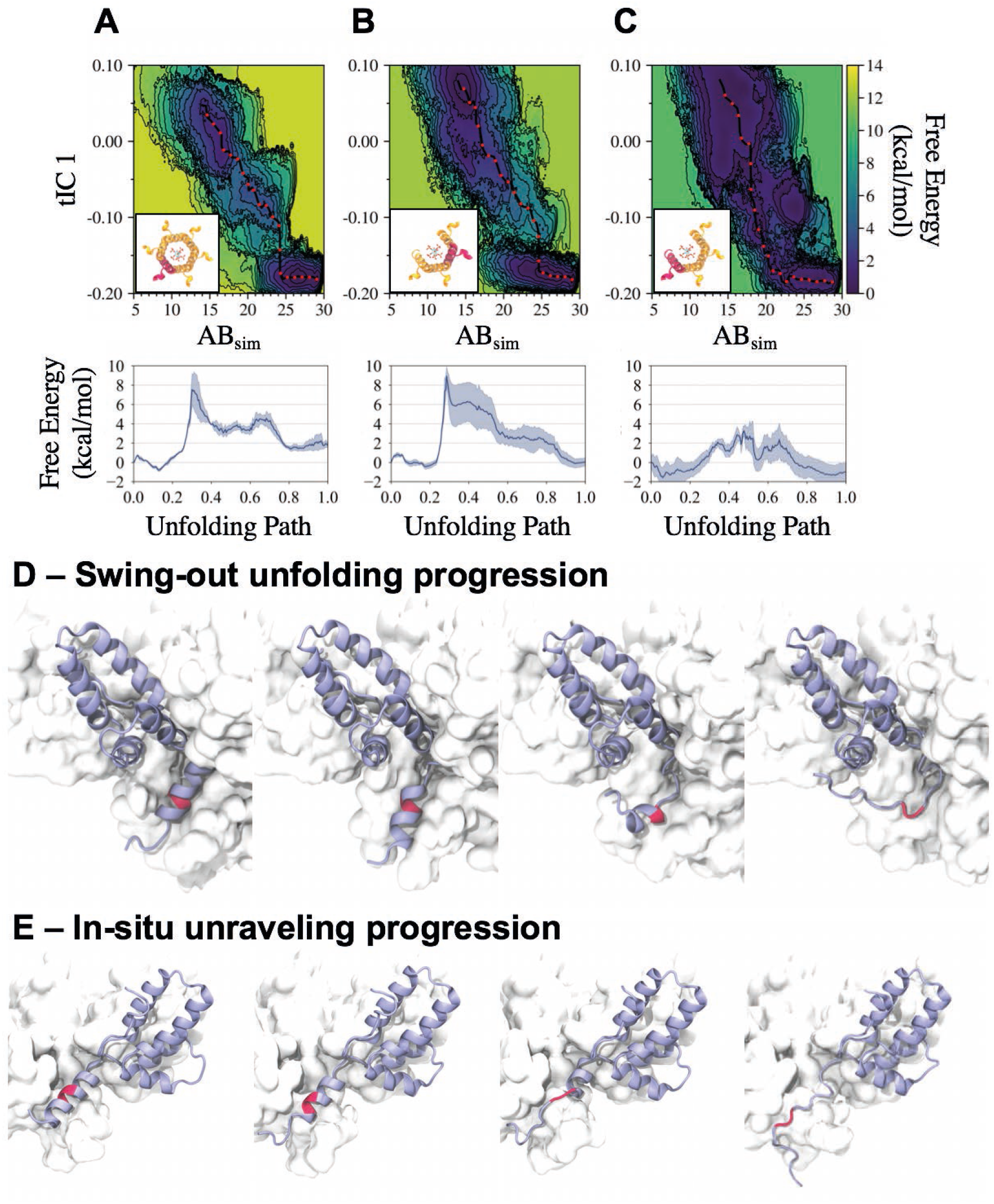
Comparison of free energy surfaces characterizing the CA-SP1 junction helix-coil transition from metadynamics simulations. The free energy is projected onto two variables – alpha-beta similarity (AB_sim_) and the first time-structure independent component (tIC_1_) for (A) a helix in a complete hexamer and helices in an incomplete hexamer missing 2 neighboring CA-SP1 monomers, where we consider (B) a helix between two neighboring helices and (C) the outer helix (with V362 and A366 exposed to solvent); the helix highlighted red in each inset represents each considered helix. Each respective minimum free energy path is depicted as a black line with red dots and quantified in the subsequent plots below. Two unfolding pathways are depicted in (D) and (E), with the former representing the primary helix-to-coil transition pathway explored in (A/B) and the latter representing the primary pathway explored in (C); the biased monomer is depicted in a purple ribbon representation while the CA-SP1 proteolytic cleavage site is depicted in red.

The two observed helix-to-coil unfolding routes are as follows. The first path is what we term the “swing-out” route, in which the helix begins unfolding below residue S368 and above residue R361 (residues V362-M367 have a propensity to stay helical) while remaining within the helical bundle, after which the helical segment escapes from the bundle and unfolds while solvated (depicted in **Fig. 5D**). The second path is what we term the *“in-situ* unraveling” route, in which the helix processively unfolds from the bottom of the helix while contacts with the adjacent helix are maintained, after which the helix completely unfolds and detaches (depicted in **Fig. 5E**). Our simulations suggest that the primary unfolding pathway for helices with two neighbors (in both complete and incomplete hexamers) is the “swing-out” route while the primary unfolding pathway for the exposed helix in incomplete hexamers is the “insitu unraveling” route.

Inositol hexakisphosphate (IP_6_) is a small molecule that binds within the central pore of immature CA-SP1 hexamers and is known to be an assembly cofactor for the immature virus (26). To test the importance of IP_6_ on the helix-coil transition, we conducted our simulations both in the presence (all data described above) and absence of IP_6_ (see Supplementary Fig. 4). Our simulations show that the absence of IP_6_ induces modest quantitative differences; the helix-to-coil transition free energy barriers are 6-7.5 kcal/mol for helices with two neighbors in complete and incomplete hexamers (compared to 8.5-9 kcal/mol computed in the presence of IP_6_). The exposed helix in the incomplete hexamer has a transition barrier of 3 kcal/mol without IP_6_ (compared to 4 kcal/mol with IP_6_). Hence, IP_6_ tends to reduce the propensity for helix-to-coil transitions, likely by stabilizing the helical bundle, but does not compensate for the increased helix-to-coil transitions expected in partial hexamers. Similarly, we find that helices in partial hexamers are more likely to explore both swing-out and in-situ unraveling transition pathways. We conclude from our simulations that the helix-to-coil transition for the exposed helix in an incomplete 6HB is more amenable to unfolding than that of complete hexamers by virtue of an alternative unfolding pathway with a free energy barrier height that is reduced by up to 5 kcal/mol.

## Discussion

### Implications for immature virus assembly

Our subtomogram averaging analysis shows structures present at the discontinuous edges of the immature Gag lattice. These structures have been derived from viruses with an inactive protease purified 44h post transfection, and subsequently purified. They therefore most likely do not represent transient intermediates, but stable end states. We observe that growth of the lattice ceases such that the lattice edge does not form at the boundary between Gag hexamers (**Fig. 6A**), but instead it forms at the boundaries between Gag dimers (**Fig. 6B, C**). In other words, incomplete Gag hexamers exist at the lattice edge, while all Gag monomers we observe are dimerized. These observations suggest that Gag lattice growth proceeds by the addition of Gag dimers.

**Figure 6.**
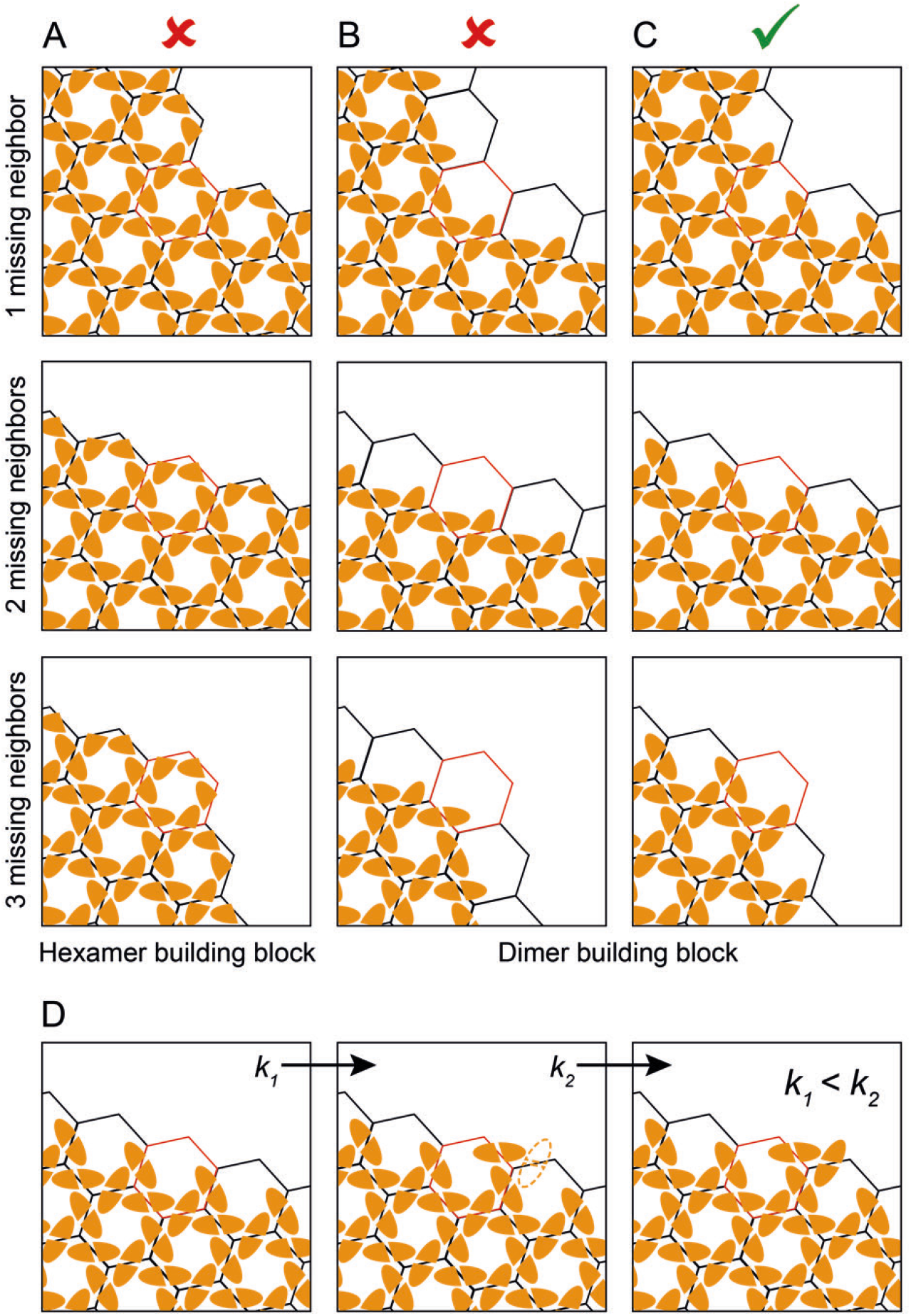
Schematic showing possible modes of Gag lattice growth and the expected structures of the lattice edges at positions with 1, 2 or 3 missing hexamers. CA_CTD_ monomers are shown as orange shapes with flat dimerization interfaces. (A) Assembly via addition of hexamers – lattice edges would consist of complete hexamers in this mode of assembly. (B, C) Assembly via addition of dimers – lattice edges would consist of complete dimers in this mode of assembly. In (B), the edges consist primarily of dimers in which one component monomer forms part of a complete hexamer giving rise to partial hexamers with 1-3 contributing monomers. In (C) the edges consist primarily of dimers in which both component monomers contribute to partial hexamers, giving rise to partial hexamers with 3-5 contributing monomers. (C) is the mode of assembly consistent with our observations. (D) Binding of a Gag dimer in which only one component monomer is part of a hexamer creates a binding site (outline) where a dimer can bind with both component monomers, as part of two hexamers. If association constant k_1_ < k_2_, assembly typically proceeds to bind this site before arresting.

Our observations are consistent with the severe assembly phenotypes of mutations in the dimer interface such as WM184,185AA (27, 28). They are also consistent with the observation that constructs in which oligomerization-promoting NC is replaced with a dimerizing leucine zipper domain are competent for assembly (29).

Each Gag dimer contributes to two CA_NTD_ trimers, and in agreement with this we observed loss of density for two CA_NTD_ trimers in our maps for each missing dimer. These results show that CA_NTD_ domains that are not fully trimerized are not packed in an ordered manner into the Gag lattice.

Our observations suggest that Gag dimers are the key assembly unit during immature virus assembly. The addition of Gag dimers to a growing lattice will continue until addition of further Gag dimers is unfavorable due to constraints of lattice geometry, or until the available Gag is depleted. This could occur such that growth typically arrests where each edge dimer has at least one monomer which is part of a complete hexamer. This would result in partial Gag hexamers at the lattice edges generally having one to three members (**Fig. 6B**). Alternatively, this could occur such that growth typically arrests where each edge dimers has both constituent monomers bound to a partial hexamer. This would result in partial hexamers with three to five members (**Fig. 6C**) and these are the classes which we observed. This observation suggests that the binding affinity of a dimer where both constituent monomers make interactions within partial bundles is higher than the binding affinity when only one monomer makes interactions, even if it completes a helical bundle. Furthermore binding of a dimer such that both constituent monomers make interactions may be favored due to the avidity effect of having two compatible binding sites. This suggests the following hierarchy of association events during assembly: if a dimer binds via one monomer it creates a site where a second dimer can bind with both monomers and the higher binding affinity makes it likely that this second binding site will be occupied (**Fig. 6D**). As a result we observe that where a site exists on the growing lattice edge that can accommodate a Gag dimer such that both component monomers join adjacent partial hexamers, then a Gag dimer is generally bound to that site.

Our findings support coarse-grained molecular dynamics that previously simulated immature Gag lattice assembly as the addition of Gag dimers through 6HB interactions (10). These simulations observed the association of Gag dimers at edges that cyclically varied between the edge cases presented in **Fig. 6B-C**. However, Gag dimers tended to favor the formation of trimer-of-dimers to maximize 6HB contacts, i.e., the state depicted in **Fig. 6C**, which further suggests that the association constant of the second binding site (*k_2_*) depicted in **Fig. 6D** is larger than that of the first binding site (*k_1_*). Partial hexamers at edges were predicted to form throughout the assembly until passivated by Gag addition and predicted to persist as lattice defects when kinetically trapped (10), consistent with our observations here.

Overall our data suggest that immature HIV-1 assembly proceeds by recruitment of Gag dimers into the growing lattice via formation of intra-hexameric interactions. The conservation of the arrangement of the CA_CTD_ structures among the retroviruses, contrasting with the variable arrangement of the CA_NTD_ (7, 30, 31), suggests to us that this assembly route is likely to be conserved.

### Implications for virus maturation

Unfolding of the CA-SP1 helical bundle appears to be the main structural determinant of the transition from the immature Gag lattice to a mature CA lattice (32). The 6HB formed by the CA-SP1 junction has been shown to exist in a helix-coil equilibrium, and this equilibrium is likely to limit access of the protease to the cleavage site within the bundle (25). Cleavage makes the transition irreversible. We observed that the CA-SP1 helical bundle was still somewhat ordered even when up to 3 Gag subunits were missing from a hexamer. As the resolution of our structures is insufficient to unambiguously resolve the structures of these partial 6HBs, we performed molecular dynamics simulations to assess their structural stability. Our simulations show that partial helical bundles can remain ordered, but that there is an increased probability of uncoiling of the CA-SP1 helix of the Gag molecule at the clockwise edge (from a top-down view) of the partial bundle. Coordination of the 6HB by IP_6_ seems to hinder uncoiling (by 1-3 kcal/mol) but is insufficient to completely inhibit uncoiling, especially in partial bundles. These observations suggest that CA-SP1 cleavage may initiate stochastically within partial hexamers at lattice edges which would then cause local disassembly of the lattice as it undergoes structural maturation. This, in turn, would destabilize hexamers immediately adjacent to the maturation event by removing the inter-hexamer interactions that are involved in both dimer and trimer formation in the immature lattice, and these hexamers would then be more likely to undergo maturation compared to those in the middle of the lattice. This would proceed towards the middle of the lattice as a ‘wave’ of maturation from one or more initiation sites at lattice edges, promoting lattice disassembly and proceeding to consume CA-SP1 inwards from that site.

## Materials and Methods

### Generation of the cryo-EM dataset

A previously published cryo-ET data set of purified, immature HIV-1 viral particles which yielded a 4.2 Å structure of the immature hexamer (9) (EMDB accession number: EMD-4017), was used as a starting point for analysis of immature Gag lattice edges. This data set consists of 74 tomograms containing 484 viruses previously used for structural determination, with an unbinned pixel size of 1.35 Å/pixel.

To ensure data completeness, we used roughly-aligned subtomogram positions from an intermediate step in the processing of the data set above (9), immediately prior to the exclusion of subtomograms based on cross-correlation value. These positions had been generated by three successive iterations of alignment and averaging of 8× binned subtomograms against an initial 6-fold symmetric reference. As subtomogram extraction positions were oversampled for the initial angular search, duplicate subtomograms that had aligned onto the same positions were removed from the data set by applying a pairwise distance criterion of 4 binned pixels (4.32 nm). The remaining positions were then visualized in UCSF Chimera using a custom plugin as described in K. Qu et al. (30), and misaligned subtomogram positions were removed by manual inspection. Misaligned positions were defined as those positions not conforming to the geometry of the hexagonal lattice, for example those that were substantially rotated out of the plane of the lattice. We did not exclude any positions based on cross-correlation coefficient (CCC), since this could result in partial hexamers being removed from the data set if they correlated less strongly to the 6-fold hexameric reference.

### Selection and preparation of lattice edge positions

The remaining 178,750 aligned subtomogram positions were then analyzed to identify the edges of the immature Gag lattice. A custom MATLAB script was used to identify every possible pattern of missing neighbors around each hexamer position in the lattice map of each virus in the data set. Using this script, we identified 62815 potential edge positions and oriented all of them to place the predicted gap in the lattice in a single direction. Noncontiguous gap classes were discarded. As the number of subtomograms missing 4 or 5 neighbors was very low, we retained only subtomograms with 1, 2 or 3 contiguous neighboring hexamers missing. This resulted in a data set of subtomogram positions containing 57134 points, which we pooled.

The coordinates of the oriented subtomogram positions for edge hexamers were scaled for use with 4× binned data. Subtomograms were extracted from 4× binned tomograms with a box edge size of 72 binned pixels, corresponding to 388.8 Å in each dimension. One iteration of angular refinement was then performed against the final 4× binned average previously generated by F. K. M. Schur et al. (9). The alignment was performed using an 8 × 2° angular search range for all Euler angles, a 32.4 Å low pass filter, C6 symmetry and a mask around the central hexamer and all six neighboring positions. The resulting subtomogram positions were used as the starting point for image-based classification. The average of the aligned subtomograms was also generated for subsequent use in wedge-masked difference map and multi-reference alignment-based classification.

### Classification by principal component analysis (PCA) of wedge-masked difference maps (WMD)

We adapted wedge-masked difference map-based subtomogram classification (hereafter referred to as WMD PCA), originally described by J. M. Heumann et al. (23) for use with subtomogram averaging scripts based on the TOM (33), AV3 (22) and Dynamo (34) packages.

To generate the missing wedge mask, 100 subtomograms were extracted from empty “noise” regions in each tomogram and normalized to a mean grey value of 0 with a variance of 1. Their amplitude spectra were calculated and averaged to generate a Fourier weight for that tomogram that describes missing information in Fourier space due to the missing wedge and the CTF – this is the wedge mask for that tomogram.

Each subtomogram was rotated into the reference frame according to the angles calculated during angular refinement. The same rotation was applied to the corresponding wedge volume. The subtomogram and normalized reference were Fourier transformed, low-pass filtered to 30 Å, multiplied by the wedge mask, and inverse Fourier transformed to generate the wedge weighted volume. A real-space mask was then applied to the weighted volumes so that only the central hexamer and its six immediate neighbors were considered for difference map calculation. The grey values under the mask were again normalized and the weighted and masked subtomogram volume was subtracted from the weighted and masked average of all rotated subtomograms in order to generate a wedge-masked difference map.

The difference map voxels under the masked region of interest were then stored as an m × n matrix, where m is the number of voxels under the mask and n is the number of subtomograms in the data set. Singular value decomposition (SVD) was performed on the matrix of difference map voxels to decompose the voxel matrix D according to the relationship D = USV^T^. The first 30 left singular vectors of the matrix were obtained from the matrix U, reshaped to match the mask, and stored as the first 30 eigenvolumes of the data set. SV^T^ was stored as this provides the corresponding eigencoefficients for use in clustering the data.

Eigenvolumes were inspected manually to identify those corresponding to structural differences between the average structure and the subtomograms, rather than those describing residual differences due to the orientation of the subtomograms relative to the missing wedge. A subset of eigenvolumes was selected. The eigencoefficients corresponding to these selected eigenvolumes were used as input for k-means clustering in MATLAB with 10 replicates and k = 30. The subtomograms in the data set were then grouped into classes based on these clusters, and the average of each class was generated. Class averages were inspected visually, and classes containing 1, 2, 3 as well as no missing hexameric neighbors around the central hexamer were identified, with some classes rotated in-plane by 1 hexamer position (i.e. 60°). The in-plane rotation angle of the subtomogram positions in these rotated classes was adjusted in order to match the configurations seen in the other classes. Multiple classes were identified as missing 1, 2 and 3 neighboring hexamers, and these classes were pooled into larger, single classes with 1, 2 and 3 missing neighboring hexamers before generation of the class averages (Supplementary Figure 2C).

### Classification by multi-reference alignment

Subtomograms were divided into equal-sized subsets according to odd and even particle number and averaged in order to produce two starting averages, each with half of the data. The odd and even half-references were then multiplied in MATLAB by masks constructed in order to down-weight 1, 2 or 3 hexamers around the central hexamer (Supplementary Fig. 3).

The original, non-multiplied references were also included to allow identification of complete hexamers. Subtomograms were then aligned against all of these artificial references for 6 iterations, with a low pass filter of 29.9 Å, a real-space mask passing the central hexamer as well as three of its contiguous neighbors in the gap direction, and a restricted in-plane angular search allowing only rotation of 60° in each direction. The reference to which each subtomogram aligned with the highest cross-correlation coefficient CCC was used to assign the class of that subtomogram. Simulated annealing was used for stochastic sampling in order to allow subtomograms to escape from local minima between alignment iterations, using a scaling factor of 0.3 with the approach described by T. Hrabe et al. (24). Class membership had converged onto stable classes by the sixth iteration, and the results of this alignment iteration were used for subsequent structure generation and analysis.

### Generation of class averages

The pooled classes from WMD PCA classification, as well as the final classes from multireference classification, were then unbinned to regenerate the class averages from 2× binned subtomograms as the final structures.

### Molecular dynamics simulations and analysis

The initial all-atom protein configuration was adopted from an atomic model (PDB 5L93); systems with missing monomers were initialized by deleting relevant monomers. Myoinositol hexakisphosphate (IP_6_)(fully deprotonated) was randomly positioned between the two rings of lysine (K290 and K359) in the central pore region and each of the six incomplete pore regions along the exterior protein interface. All proteins were solvated by water and 150 mM NaCl in a rhombic dodecahedron simulation domain large enough to contain a 1.5 nm layer of water perpendicular to each exterior protein interface. Energy minimization was performed using steepest descent until the maximum force was less than 1000 kJ/mol/nm. Equilibration was performed with harmonic restraints (using a 1000 kJ/mol/nm^2^ spring constant) on each heavy atom throughout the protein for 10 ns in the constant *NVT* ensemble using stochastic velocity rescaling (35) at 310 K and a damping time of 0.1 ps. Restraints were then removed and each system was allowed to equilibrate for 100 ns in the constant *NPT* ensemble using a Nosé-Hoover thermostat (36) (2 ps damping time) and a Parrinello-Rahman barostat (37) (10 ps damping time) at 310 K and 1 bar.

Two types of production runs were performed. Simulations to characterize the flexibility in protein structure were performed over 300 ns with configurations saved every 20 ps. Rigid structural alignment of the protein complex in each frame was performed with the CTD domains of the interior CA-SP1 hexamer in reference to the atomic model. The atomic model was also used as reference to compute RMSDs after alignment.

Simulations to characterize free energies were performed over 850 ns and used the following two coordinates. The first is the alpha-beta similarity (AB_sim_) which is given by:

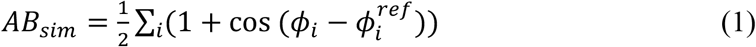

where *i* denotes an index over the considered dihedrals and residues, *ϕ_i_* is the dihedral angle, and 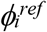 is the reference dihedral angle. Here, we include the phi and psi angles of the CA-SP1 junction (Gag residues 356-370) and set 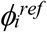 to −60 degrees, such that when the junction is completely α-helical, AB_sim_ is 30.

The second are components identified from tICA (38), which is a technique used to identify linear projections of data (i.e., linear combinations of features) that capture the slowest varying motions by maximizing the autocorrelation function; here, we use cos(*ϕ_i_*) as our feature set and a lag time (τ) of 35 ns to construct a covariance matrix:

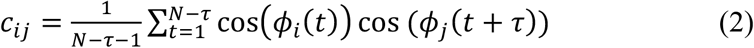

Each tIC is each eigenvector identified by solving the generalized eigenvalue problem. Here, we consider the first tIC (i.e., the slowest mode or the eigenvector corresponding to the largest eigenvalue) as our second free energy coordinate.

The WT-MetaD algorithm (39) was used with a Gaussian bias (using a height and width of 0.5 kJ/mol and 0.1) deposited every 1 ps along the AB_sim_ coordinate with a bias factor of 25 *k_B_T*. The resultant free energy surfaces were projected onto different coordinates using the Tiwary-Parrinello reweighting algorithm (40). The MSMBuilder python library (41) was used to perform tICA on a separate MD trajectory of a peptide representing the CA-SP1 junction. The second tIC was also considered as an additional coordinate for dimensional reduction of the underlying free energy surface (see Supplementary Fig. 5).

All simulations were prepared and simulated with GROMACS 2016 (42) using the CHARMM36m forcefield (43). A timestep of 2 fs was used in all simulations with hydrogen-containing bonds constrained using the LINCS algorithm (44). Metadynamics simulations were performed using the PLUMED 2.4 plugin (45). Table 1 summarizes relevant statistics for each simulated system.

**Table 1.**
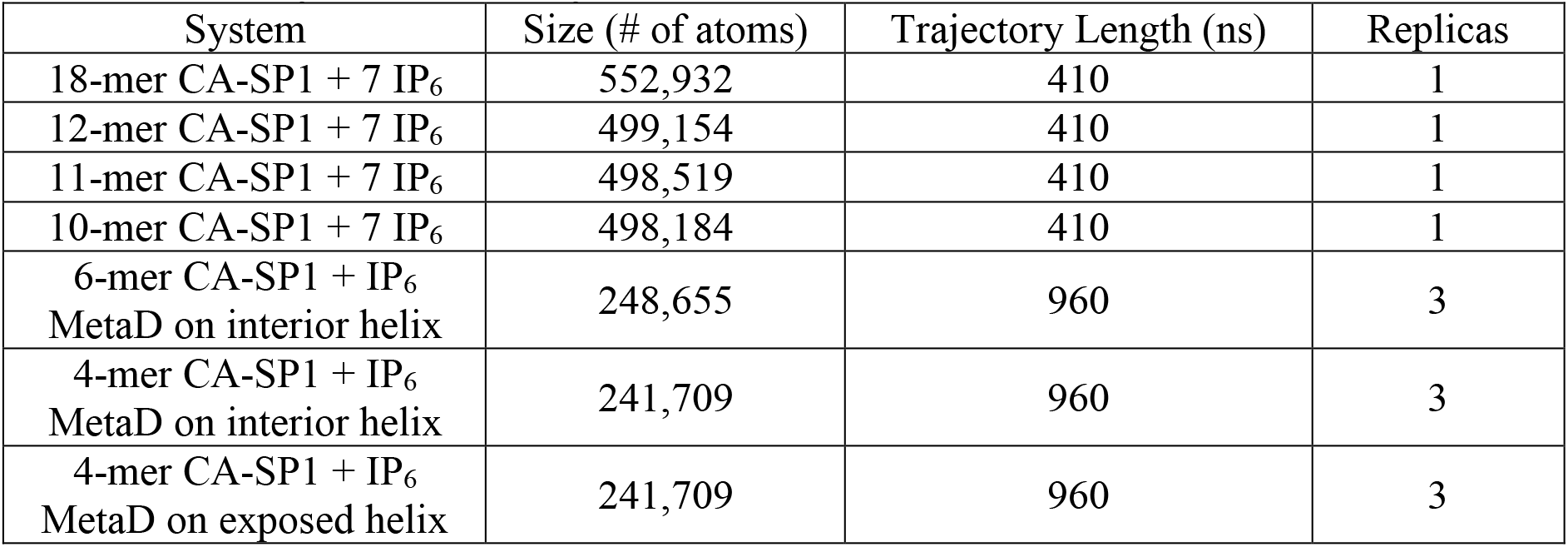
Summary of molecular dynamics simulations details.

| System | Size (# of atoms) | Trajectory Length (ns) | Replicas |
| --- | --- | --- | --- |
| 18-mer CA-SP1 + 7 IP<sub>6</sub> | 552,932 | 410 | 1 |
| 12-mer CA-SP1 + 7 IP<sub>6</sub> | 499,154 | 410 | 1 |
| 11-mer CA-SP1 + 7 IP<sub>6</sub> | 498,519 | 410 | 1 |
| 10-mer CA-SP1 + 7 IP<sub>6</sub> | 498,184 | 410 | 1 |
| 6-mer CA-SP1 + IP<sub>6</sub><br>MetaD on interior helix | 248,655 | 960 | 3 |
| 4-mer CA-SP1 + IP<sub>6</sub><br>MetaD on interior helix | 241,709 | 960 | 3 |
| 4-mer CA-SP1 + IP<sub>6</sub><br>MetaD on exposed helix | 241,709 | 960 | 3 |

### Model fitting and lattice maps

An atomic model derived from a high-resolution cryo-EM structure of the immature Gag CA domain hexamer (PDB 5L93) was used for all model fitting into the cryo-EM density maps (9). The coordinates for the CA_NTD_ and CA_CTD_ from this model were fit as rigid bodies into each corresponding position in the density maps using UCSF Chimera (46). This was done for each of the density maps containing hexamers with one, two or three missing CA protomers from the central hexamer position. The atomic models shown in positions where there was no CA density (**Fig. 2**) were positioned for illustration purposes using the corresponding density map of the average complete hexamer from this dataset (9).

Lattice maps showing the positions and orientations of aligned subtomograms within the coordinate system of the original tomograms were plotted in UCSF Chimera using a custom plugin (30).

## Acknowledgements

We acknowledge Kun Qu for providing advice and assistance in data processing and classification of subtomogram data. We also acknowledge Jake Grimmett and Toby Darling from MRC LMB Scientific Computing for providing technical support. A.J.P. acknowledges support from the Ruth L. Kirschstein National Research Service Award Postdoctoral Fellowship by the National Institutes of Health (F32-AI150477). G.A.V. acknowledges support from the National Institute of Allergy and Infectious Diseases of the National Institutes of Health (R01-AI150492). This work used computational resources provided by the Extreme Science and Engineering Discovery Environment (XSEDE), which is supported by National Science Foundation grant number ACI-1548562. J.A.G.B acknowledges funding from the European Molecular Biology Laboratory (EMBL) and Medical Research Council Grant MC_UP_1201/16.

---

**Supplementary Figure 1.**
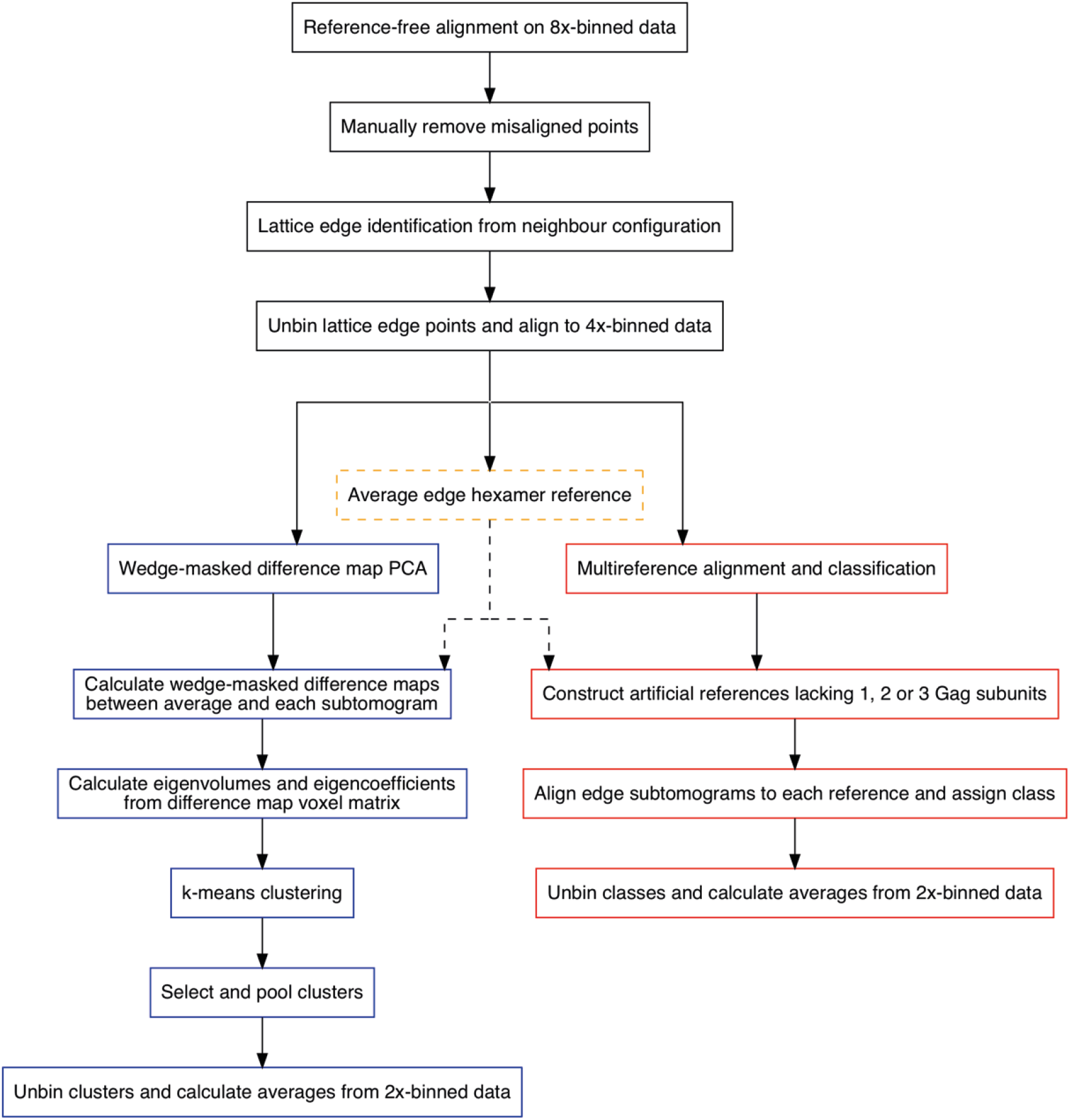
Subtomogram alignment and classification workflow used to determine Gag lattice edge structures. A dataset of 8× binned subtomograms from (9), which had previously been aligned reference-free as described in Materials and Methods, was used as a starting point for manual removal of misaligned points (black boxes). An initial geometric identification and re-orientation of hexamers along lattice edges from the configuration of neighboring subtomograms was then performed, followed by extraction of subtomograms centered on the identified coordinates from 4× binned data (black boxes). An initial average reference containing all identified edge hexamers was constructed using 4× binned data as described in Materials and Methods (dashed yellow box). This reference was to calculate wedge-masked difference maps against each subtomogram (blue boxes), and separately also to construct synthetic references for multireference alignment and classification (red boxes). These two classification approaches were carried out completely independently on the same input data.

**Supplementary Figure 2.**
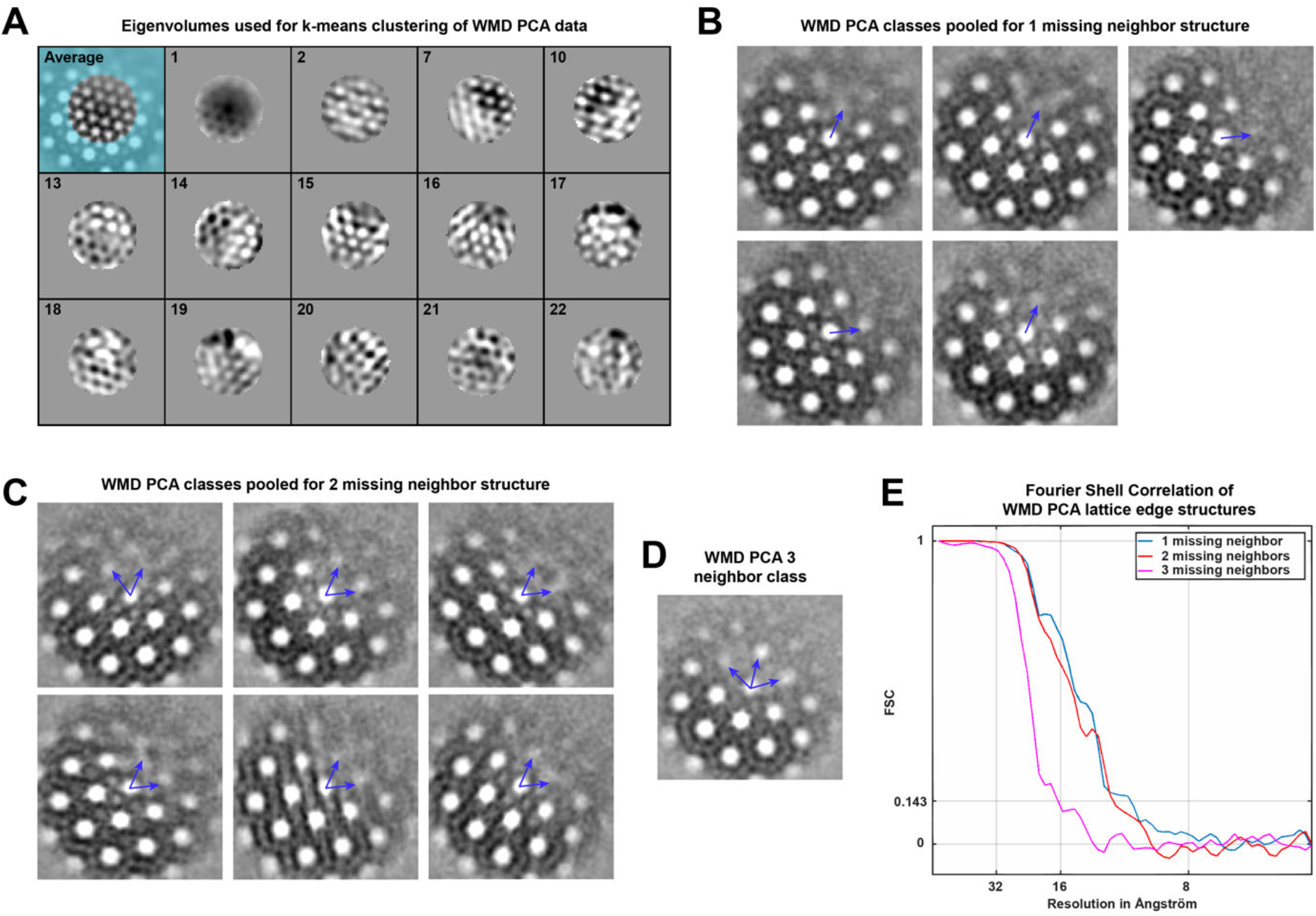
Image processing details for WMD PCA classification of lattice edge hexamers. (A) Central XY slices through eigenvolumes selected as the principal components defining the lower-dimensional space onto which subtomograms were projected for classification, labelled with corresponding principal component number. The top left panel shows the average structure with an overlaid binary mask defining the voxels used for difference map calculation, with cyan regions not considered. (B) Classes from k-means clustering based on wedge-masked difference maps, corresponding to hexamers with 1 missing neighbor. Two classes were rotated by 60° relative to the other classes, corresponding to inaccuracies in the initial geometric orientation of the missing neighbor position (positions denoted by blue arrows extending from the central hexamer, see Materials and Methods). (C) As in B, for hexamers missing 2 neighbors. (D) As in B and C, for the single class of hexamers missing 3 neighbors. (E) Fourier shell correlation (FSC) curves between the odd and even half-datasets for each partial hexamer structure after further alignment with 2× binned data (see Materials and Methods).

**Supplementary Figure 3.**
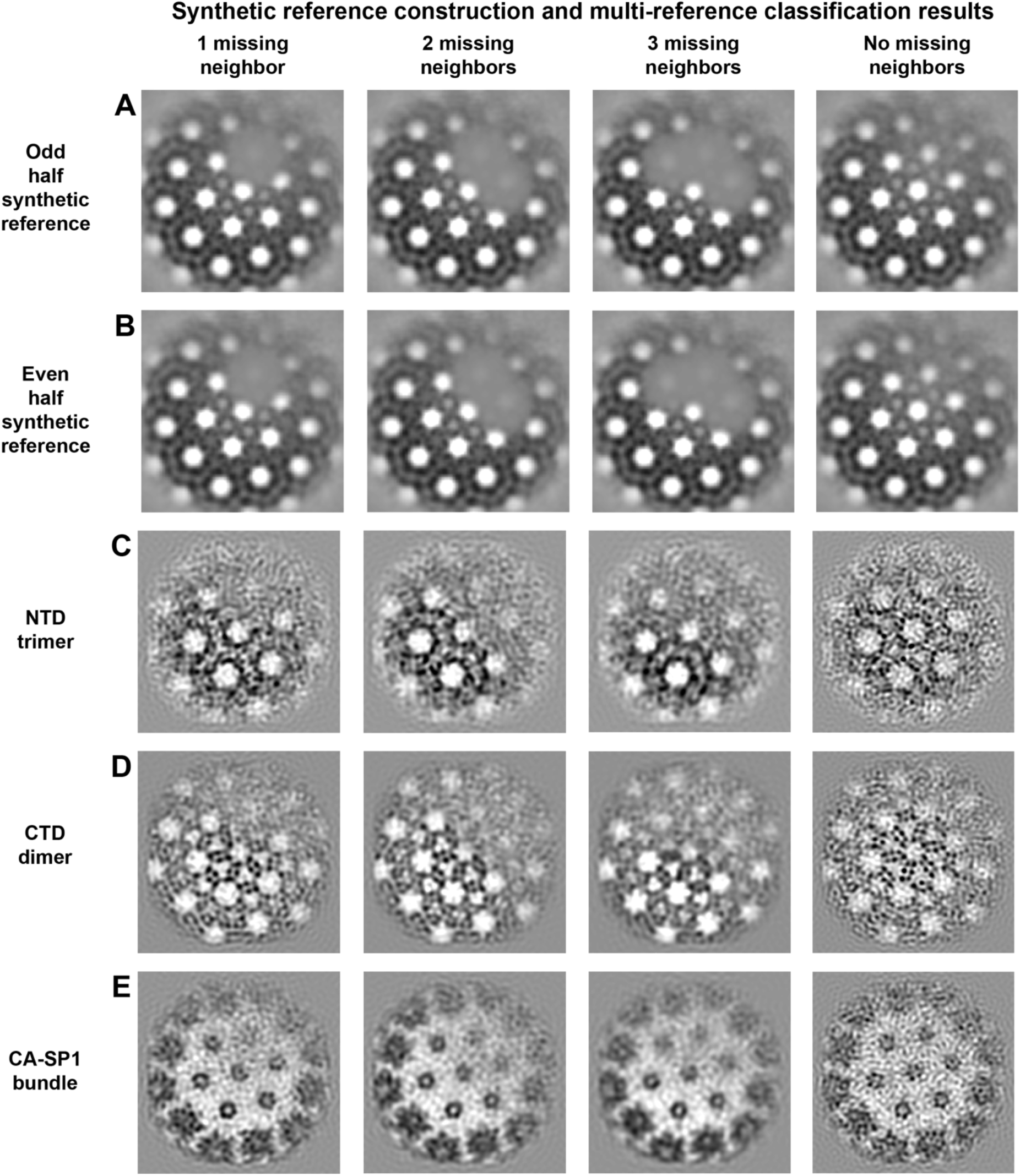
Synthetic references and subtomogram alignment results from multi-reference subtomogram alignment and classification of Gag lattice edge hexamers missing different numbers of neighbors. Synthetic references (A-B) were constructed by down-weighting density corresponding to individual hexamer positions by masking as described in Materials and Methods. Panels (A) corresponds to the synthetic references constructed using the odd half-dataset average, and panel (B) correspond to those constructed using the even half-dataset average. (C) Orthoslices through the CA_NTD_, (D) through the CA_CTD_ and (E) through the CA-SP1 helical bundle layers of the resulting final class averages from multi-reference alignment and classification are also shown for the classes corresponding to positions with 0, 1, 2 and 3 missing neighboring hexamers.

**Supplementary Figure 4.**
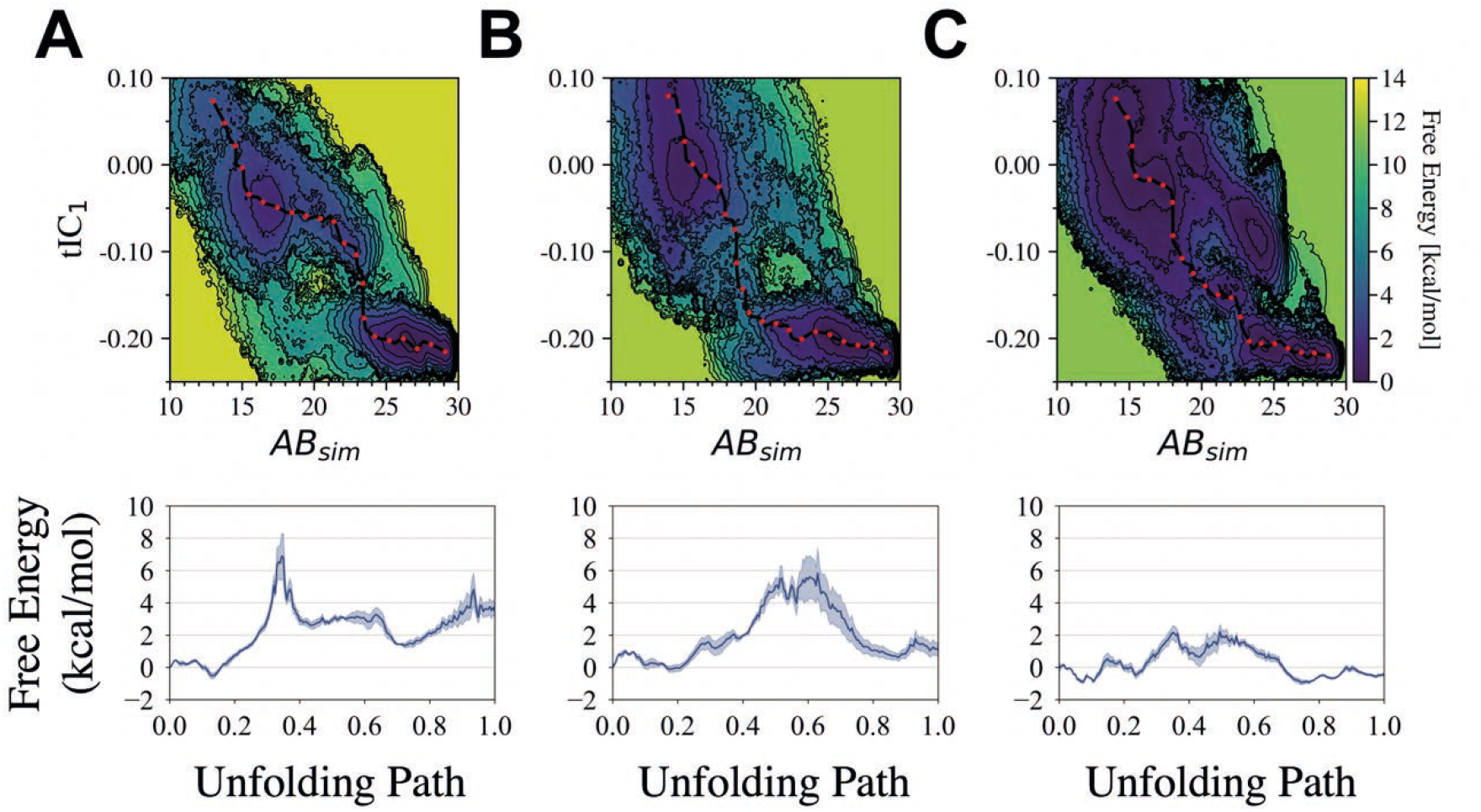
Comparison of free energy surfaces projected onto the alpha-beta similarity (AB_sim_) and first time-structure independent component (tIC_1_) for 6HBs in the absence of IP_6_. We compare (A) a helix in a complete hexamer to (B, C) helices in an incomplete hexamer missing 2 neighboring CA-SP1 monomers, where (B) is a helix between two neighboring helices and (C) is the outer helix (with V362 and A366 exposed to solvent). Each respective minimum free energy path is depicted as a black line with red dots and quantified in each of the bottom plots.

**Supplementary Figure 5.**
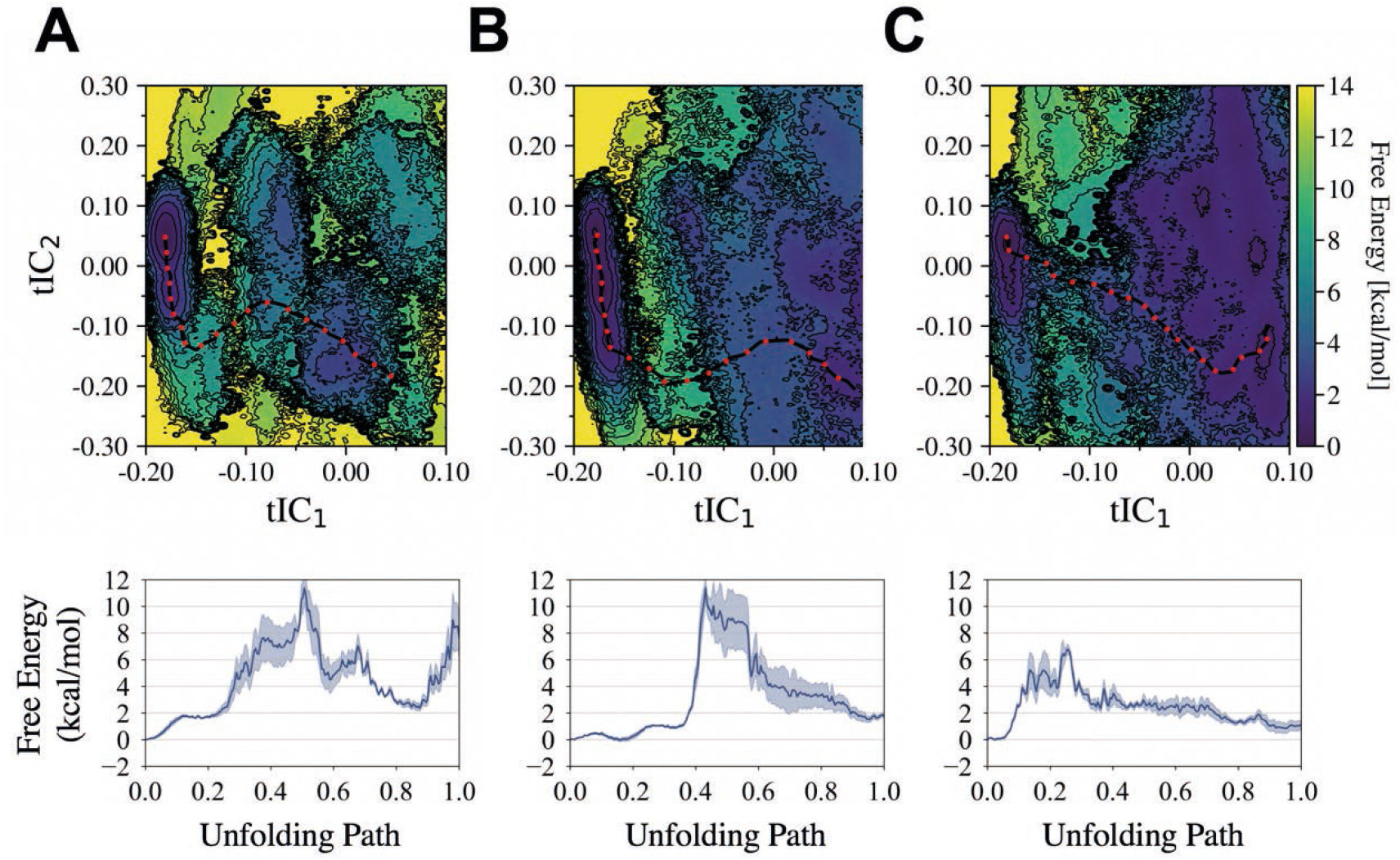
An alternate comparison of free energy surfaces projected onto the first (tIC_1_) and second (tIC_2_) time-structure independent components. We compare (A) a helix in a complete hexamer to (B, C) helices in an incomplete hexamer missing 2 neighboring CA-SP1 monomers, where (B) is a helix between two neighboring helices and (C) is the outer helix (with V362 and A366 exposed to solvent). Each respective minimum free energy path is depicted as a black line with red dots and quantified in each of the bottom plots. In each case, the 6HB is coordinated by IP_6_.

